# Multi-omics network-based functional annotation of unknown Arabidopsis genes

**DOI:** 10.1101/2021.06.17.448819

**Authors:** Thomas Depuydt, Klaas Vandepoele

**Affiliations:** Department of Plant Biotechnology and Bioinformatics, Ghent University, Technologiepark 71, 9052 Ghent, Belgium; VIB Center for Plant Systems Biology, Technologiepark 71, 9052 Ghent, Belgium; Bioinformatics Institute Ghent, Ghent University, Technologiepark 71, 9052 Ghent, Belgium

## Abstract

Unraveling gene functions is pivotal to understand the signaling cascades controlling plant development and stress responses. Given that experimental profiling is costly and labor intensive, the need for high-confidence computational annotations is evident. In contrast to detailed gene-specific functional information, transcriptomics data is widely available in both model and crop species. Here, we developed a novel automated function prediction (AFP) algorithm, leveraging complementary information present in multiple expression datasets through the analysis of study-specific gene co-expression networks. Benchmarking the prediction performance on recently characterized *Arabidopsis thaliana* genes, we showed that our method outperforms state-of-the-art expression-based approaches. Next, we predicted biological process annotations for known (n=15,790) and unknown (n=11,865) genes in *A. thaliana* and validated our predictions using experimental protein-DNA and protein-protein interaction data (covering >220 thousand interactions in total), obtaining a set of high-confidence functional annotations. 5,054 (42.6%) unknown genes were assigned at least one validated annotation, and 3,408 (53.0%) genes with only computational annotations gained at least one novel validated function. These omics-supported functional annotations shed light on a variety of developmental processes and molecular responses, such as flower and root development, defense responses to fungi and bacteria, and phytohormone signaling, and help alleviate the knowledge gap of biological process annotations in Arabidopsis. An in-depth analysis of two context-specific networks, modeling seed development and response to water deprivation, shows how previously uncharacterized genes function within the respective networks. Moreover, our AFP approach can be applied in future studies to facilitate gene discovery for crop improvement.

**Significance statement:** For the majority of plant genes, it is unknown in which processes they are involved. Using a multi-omics approach, leveraging transcriptome, protein-DNA and protein-protein interaction data, we functionally annotated 42.6% of unknown *Arabidopsis thaliana* genes, providing insight into a variety of developmental processes and molecular responses, as well as a resource of annotations which can be explored by the community to facilitate future research.

## Introduction

Over the last two decades, more than 600 plant genomes have been sequenced (NCBI Resource Coordinators 2018). Yet, little is known about how the majority of plant genes function and in which processes they are involved (Rhee and Mutwil 2014). These gene functions can be formally described using controlled vocabularies, such as Gene Ontology (GO; Ashburner et al. 2000), Plant Ontology and Plant Trait Ontology (Cooper et al. 2018) and MapMan (Thimm et al. 2004). Of these, GO is the most widely used and consists of three domains representing three aspects of gene/protein function: Cellular Component (CC), Molecular Function (MF) and Biological Process (BP). The latter describes which role a gene fulfills in the biological system, rather than how (MF) or where (CC) it executes that function. Additionally, a gene-GO-term association (shortly, a GO annotation) is explicitly coupled to an evidence term, categorized into experimental, curated and computational, and should be considered when interpreting GO annotations.

While the number of sequenced genomes has increased eightfold since 2014 (83 to 646; Figure S1a), the paucity of functionally annotated genes illustrated by Rhee and Mutwil still persists seven years later (Figure S1b). For example, it was estimated that in rice, no more than 1% of protein coding genes had been described experimentally, and to date, this fraction has not exceeded 2%. Furthermore, the model organism *Arabidopsis thaliana* remains one of the few plant species with a substantial number of experimentally characterized gene functions (BP+MF: 13,638 out of 27,655 protein coding genes), with community annotation efforts overseen by The Arabidopsis Information Resource (TAIR; Berardini et al. 2015). However, due to the laborious nature of experimental profiling and lack of sequence similarity to characterized genes, still 32.4% (8,960) of *A. thaliana* genes remain without any BP or MF annotation, regardless of the evidence term. Moreover, for other plant species the knowledge gap is even greater, with some species lacking any annotations supported by experimental evidence (Figure S1b).

To alleviate this knowledge gap, automated function prediction (AFP) methods have been developed and applied to plant genomes (Bolger et al. 2018). Classically, sequence similarity and/or domain presence methods, projecting annotations from experimentally characterized genes, are applied to annotate plant genomes. However, such sequence-based approaches are generally more suited towards inference of molecular function, and plant genomes contain many genes lacking sequence similarity to experimentally characterized genes, limiting the scope of sequence-based methods (Vandepoele and Van de Peer 2005; Rhee and Mutwil 2014).

An alternative manner to infer gene function, is through the analysis of gene co-function networks, which describe gene-gene connections (edges) based on functional coherence, e.g. shared presence in a protein complex or taking part in the same metabolic pathway. For *A. thaliana*, various gene co-function network databases exist, with examples such as AraNet (Lee et al. 2015) and STRING (Szklarczyk et al. 2019) integrating different types of functional genomics data. Furthermore, recent efforts have combined a selection of co-function networks into an ensemble network (Hansen et al. 2018). While some databases include gene expression data for a selection of plant species (e.g. PlaNet; Proost and Mutwil 2017), it is possible that the species of interest is missing. Conversely, stand-alone network inference and/or function prediction tools have the potential to reach beyond popular species included in public databases.

Large-scale benchmark studies have compared function prediction performance of different AFP methods across multiple species, including *A. thaliana* (Zhou et al. 2019). There, most top-scoring methods apply machine learning and genomic features to transfer annotations to unknown genes (Mahood et al. 2020). Some methods include additional features such as protein interaction, protein structure and gene expression. However, given the aforementioned limitations of sequence-based approaches, combined with the scarcity of experimental functional genomics data in most plant species, expression-based AFP methods exploiting widely available data from transcriptomics experiments (Figure S1b) offer a valid alternative to sequence-based annotation pipelines in plants.

Interpretation of large-scale transcriptome datasets is often facilitated by means of genome-wide co-expression analysis. Co-expression relationships between individual genes are modeled into a fully or partially connected gene network, with subsequent analysis revolving around the observation that genes involved in the same biological process tend to cluster together in the network. Therefore, gene function can be inferred from the function of its neighboring genes, which is commonly referred to as the guilt-by-association (GBA) principle, and has proven successful in plants (Heyndrickx and Vandepoele 2012; Rhee and Mutwil 2014; Rao and Dixon 2019). For example, in a study describing the transcriptome dynamics during the seed-to-seedling transition in *A. thaliana*, Silva and colleagues predicted functions for uncharacterized genes in seedling establishment (Silva et al. 2016). Likewise, a comparative transcriptomics study identified conserved lignin biosynthesis modules in angiosperms, thereby predicting novel players in cell wall biosynthesis in the grass model species *Brachypodium distachyon* (Sibout et al. 2017). These examples highlight the potential of large-scale co-expression analysis for gene function hypothesis generation in plants.

In the current study, we aimed to reduce the knowledge gap of biological process annotations in *A. thaliana* by applying co-expression analysis using a large-scale expression compendium to link biological process annotations to unknown genes. For this, a novel expression-based AFP approach was developed and, using a challenging gene annotation test set, it was shown to outperform state-of-the-art expression-based methods. Moreover, we leveraged the abundance of experimental protein-DNA interaction (PDI) and protein-protein interaction (PPI) data available for *A. thaliana* and validated our expression-based predictions in a multi-omics setting, shedding light on a variety of developmental processes and molecular responses. An in-depth look at two context-specific networks, modeling seed development and response to water deprivation, shows how previously unknown genes function within the respective networks. Additionally, as a proof of concept, we show that our expression-based AFP approach can be applied in crop species, complementing purely sequence-based annotation pipelines.

## Results

### A novel gene function prediction approach outperforms state-of-the-art expression-based methods

To compile a diverse set of expression atlases capturing a variety of plant organs, developmental stages, conditions and biological processes for *A. thaliana*, we manually curated a shortlist of RNA-Seq studies using Curse (Vaneechoutte and Vandepoele 2019), comprising a total of 18 studies (Table 1). All samples were processed using Prose (Vaneechoutte and Vandepoele 2019), to obtain aggregated gene-level TPM values. The complete dataset, referred to as the full compendium, comprises 860 samples and contains 26,267 expressed genes (TPM > 2). All studies, referred to as (individual) atlases, contain uniquely expressed genes (i.e. genes only expressed in the respective atlas), highlighting the complementary nature of the different datasets.

**Table 1.**
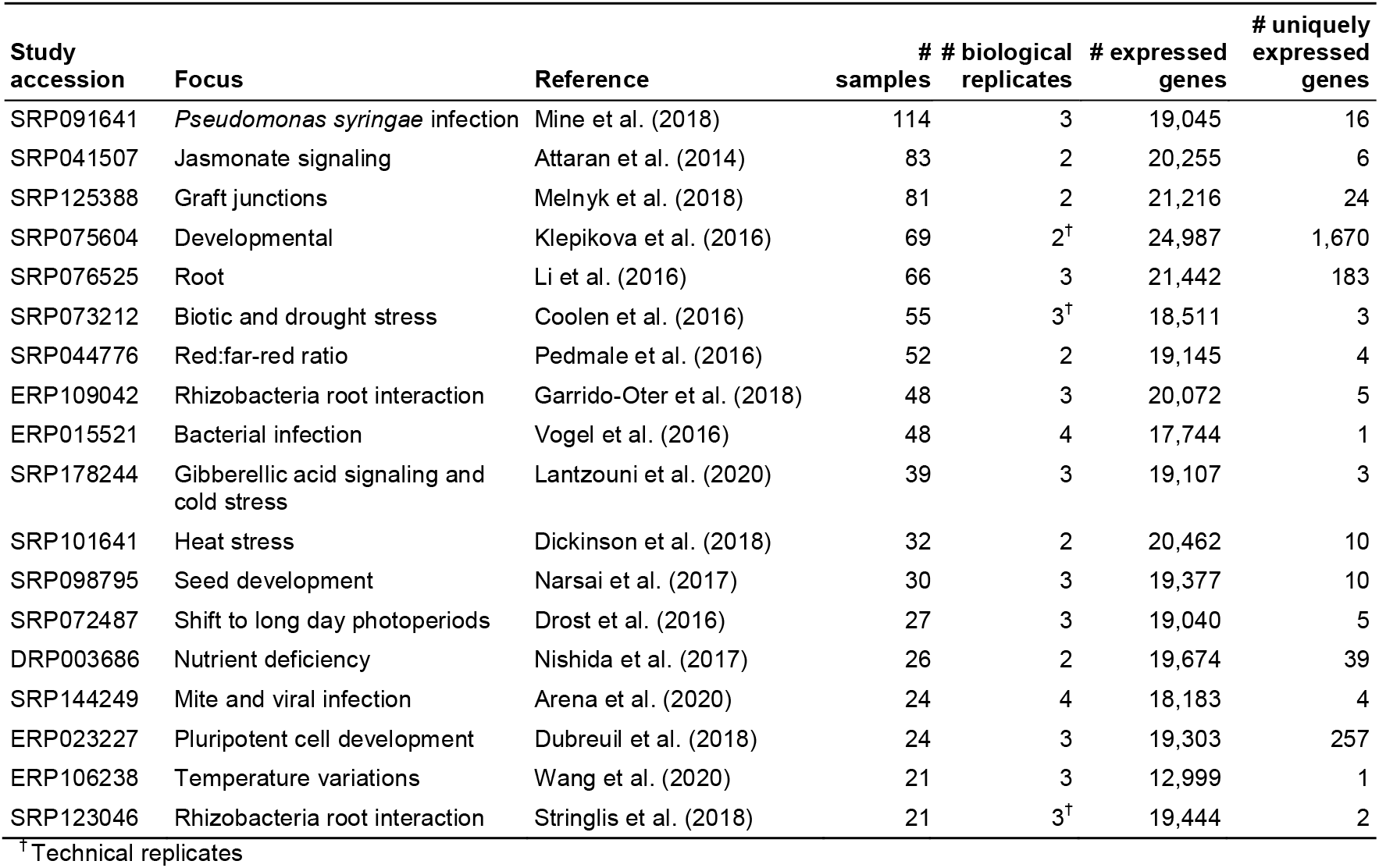
Overview of the studies included in the gene expression compendium. All 18 studies are publicly available on SRA (queried 09/04/2020) and were manually curated using Curse.

| Study accession | Focus | Reference | # samples | # biological replicates | # expressed genes | # uniquely expressed genes |
| --- | --- | --- | --- | --- | --- | --- |
| SRP091641 | *Pseudomonas syringae* infection | Mine et al. (2018) | 114 | 3 | 19,045 | 16 |
| SRP041507 | Jasmonate signaling | Attaran et al. (2014) | 83 | 2 | 20,255 | 6 |
| SRP125388 | Graft junctions | Melnyk et al. (2018) | 81 | 2 | 21,216 | 24 |
| SRP075604 | Developmental | Klepikova et al. (2016) | 69 | 2<sup>†</sup> | 24,987 | 1,670 |
| SRP076525 | Root | Li et al. (2016) | 66 | 3 | 21,442 | 183 |
| SRP073212 | Biotic and drought stress | Coolen et al. (2016) | 55 | 3<sup>†</sup> | 18,511 | 3 |
| SRP044776 | Red:far-red ratio | Pedmale et al. (2016) | 52 | 2 | 19,145 | 4 |
| ERP109042 | Rhizobacteria root interaction | Garrido-Oter et al. (2018) | 48 | 3 | 20,072 | 5 |
| ERP015521 | Bacterial infection | Vogel et al. (2016) | 48 | 4 | 17,744 | 1 |
| SRP178244 | Gibberellic acid signaling and cold stress | Lantzouni et al. (2020) | 39 | 3 | 19,107 | 3 |
| SRP101641 | Heat stress | Dickinson et al. (2018) | 32 | 2 | 20,462 | 10 |
| SRP098795 | Seed development | Narsai et al. (2017) | 30 | 3 | 19,377 | 10 |
| SRP072487 | Shift to long day photoperiods | Drost et al. (2016) | 27 | 3 | 19,040 | 5 |
| DRP003686 | Nutrient deficiency | Nishida et al. (2017) | 26 | 2 | 19,674 | 39 |
| SRP144249 | Mite and viral infection | Arena et al. (2020) | 24 | 4 | 18,183 | 4 |
| ERP023227 | Pluripotent cell development | Dubreuil et al. (2018) | 24 | 3 | 19,303 | 257 |
| ERP106238 | Temperature variations | Wang et al. (2020) | 21 | 3 | 12,999 | 1 |
| SRP123046 | Rhizobacteria root interaction | Stringlis et al. (2018) | 21 | 3<sup>†</sup> | 19,444 | 2 |

<sup>†</sup> Technical replicates

To fully exploit the wealth of available gene expression data, we developed a novel integrative network propagation AFP method that leverages the specificity of individual gene expression datasets. We conducted a benchmark study to explore and compare its performance, usability and computational requirements against a selection of state-of-the-art expression-based AFP methods. As a functional classification scheme, the Biological Process (BP) aspect from GO is used, filtered for annotations with experimental and curated evidence codes (limited to 500 GO terms annotated to 8,265 genes; see Experimental Procedures). In line with the CAFA benchmark studies (Zhou et al. 2019), a temporal hold-out (THO) validation scheme was adopted, allowing for validation on previously unknown genes (499 genes and 709 GO BP terms). All methods take as input one or more gene expression datasets and GO annotation data from the species of interest. An exception is the BLAST scoring method, included to compare the benchmark findings to CAFA, which uses sequence similarity to project GO annotations in the GOA database (Huntley et al. 2015) from BLAST hits to the (un)known query genes (Radivojac et al. 2013). For methods using only a single input dataset, a high-quality developmental atlas was selected, covering a multitude of organs such as root, hypocotyl, cotyledons and flowers (SRP075604, Klepikova et al. 2016), which translates into the highest number of expressed genes (90.4% of protein coding genes), compared to other datasets (Table 1).

Figure 1 shows a summary of the benchmark results, with green bars showing results for single datasets. Following the guidelines from Plyusnin et al. (2019), we report two main metrics to estimate overall predictor performance: (1) the gene-centric (GC) weighted Jaccard index (simGIC), defined as the intersection over the union of predictions and annotations (Pesquita et al. 2007; Figure 1a) and (2) the GC weighted F1-score (i.e. the harmonic mean of the recall and precision; Figure 1b). Here, weighted indicates that the specificity of individual predictions is considered, so that more detailed predictions (i.e. specific GO BP terms) have an increased impact on the performance metric (see Experimental Procedures). In addition, the GC weighted recall and precision are reported (Figure 1c) to uncover the balance between false negative and false positive predictions. The recall, or true positive rate, reflects the fraction of true predictions to true annotations, while the precision reflects the fraction of true predictions to all predictions. A detailed description of these and additional metrics shown in Figure 1d-g is provided in the Experimental Procedures section.

**Figure 1.**
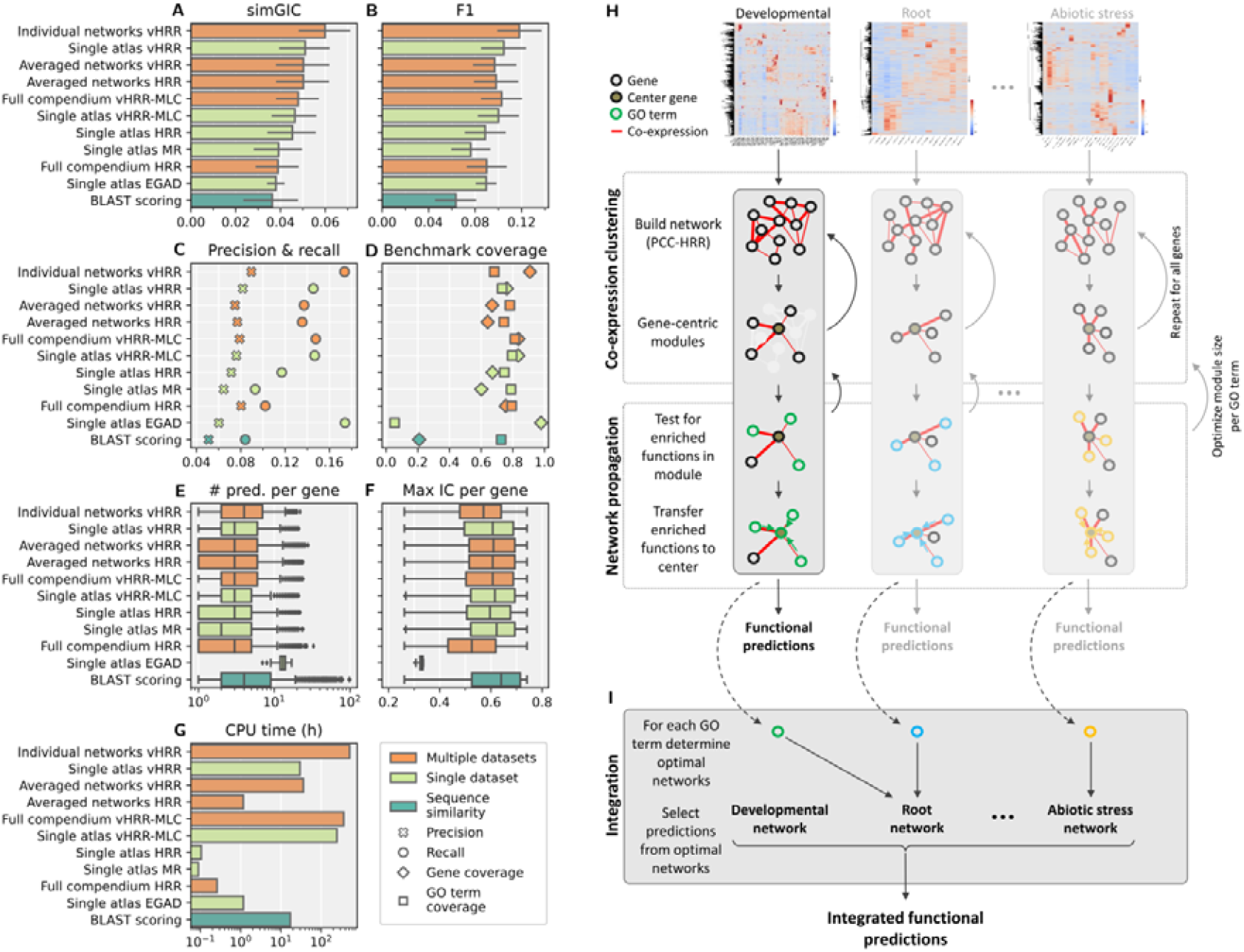
Prediction performance benchmark of expression-based AFP methods. **(A-G)** Methods are sorted based on the simGIC score and color coded based on their category. **(A-C)** Weighted gene-centric performance measures. Error bars denote the 95% empirical bootstrap confidence intervals (n=10,000). **(D)** Coverage of the temporal hold-out test set genes and GO terms. **(E)** Distribution of the mean number of predicted GO terms per gene, after removal of parental terms. **(F)** Granularity of functional inference, measured as the distribution of information content (IC) values of the most specific GO term predicted for each gene. **(G)** CPU time in log scale.Cartoon of the “Single atlas vHRR” protocol (single highlighted column). A full description is provided in the Experimental Procedures section. PCC: Pearson correlation coefficient, HRR: highest reciprocal ranks. **(H-I)** Cartoon of the “Individual networks vHRR” protocol. A full description is provided in the Experimental Procedures section.

The novel AFP method presented here uses gene expression data to build gene co-expression networks and applies the GBA principle by means of enrichment analysis (Figure 1h). As described in the Experimental Procedures section, the Single atlas HRR approach transforms the initial gene co-expression network, where edge weights represent pairwise Pearson correlation coefficients (PCC), to a fully connected highest reciprocal rank (HRR) network. This methodology was previously reported to outperform other strategies (Liesecke et al. 2018). However, Liesecke et al. (2018) did not consider mutual rank (MR), a popular alternative to HRR applied among others by the ATTEDII database (Obayashi et al. 2018). Figure 1a-c shows that HRR outperforms MR in our THO validation setup (simGIC: MR: 0.039, HRR: 0.045, relative increase of 15.6%). For these methods, the co-expression neighborhood size is optimized globally to maximize the training F1-score. A variation where the module size is optimized for each GO term individually, named variable-HRR (vHRR), further improves prediction quality by 12.7% (simGIC: 0.051), albeit at the cost of an increased computational time (two orders of magnitude; Figure 1g).

Considering each GO term individually not only allows to vary module sizes, but also to build term-specific networks. The Metric Learning for Co-expression (MLC) method was applied to learn sample weights per term, from which term-specific co-expression networks were constructed (Makrodimitris et al. 2020). However, evaluation results indicate that this approach does not improve prediction quality (simGIC: vHRR: 0.051, vHRR-MLC: 0.046), while further increasing the computational requirements (Figure 1a-c,f). Therefore, the inclusion of MLC is deemed not advantageous when only considering a single expression dataset.

Extending ‘Guilt-by-Association’ by Degree (EGAD) is a network propagation method which implements a neighbor voting algorithm that considers every gene in the network to calculate posterior probabilities for all possible gene-GO combinations (Ballouz et al. 2017). Our benchmark setup requires a well-defined set of predictions, and therefore an additional step was added to optimize the posterior probability threshold (see Experimental Procedures). The resulting predictions were evaluated and overall performance measures are included in Figure 1 (a-f). Most notable is that EGAD, at the optimal prediction score threshold, predicts only a small fraction of evaluation GO terms (38 out of 709) for almost all evaluation genes (488 out of 499; Figure 1d). Additionally, predicted functions are less specific compared to other methods (Figure 1f). Combined with suboptimal simGIC and F1-scores (0.038 and 0.090, respectively), both at the optimized posterior probability threshold as well as less stringent thresholds (Figure 1a,b; Figure S2), our benchmark indicates that EGAD is not ideal for functional prediction of unknown genes in Arabidopsis. However, it should be noted that the EGAD package includes additional functionalities not tested here. Figure 1a,b show performance metrics for the BLAST scoring method and indicate that it performs subpar compared to all other methods, which is in line with the findings from CAFA (Zhou et al. 2019).

Taken together, our newly presented expression-based AFP approach, which applies GO enrichment analysis and optimized gene neighborhoods per GO term, outperforms baseline sequence similarity and state-of-the-art expression-based methods.

### Leveraging multiple expression atlases improves gene function prediction

Next to designing an improved network-based gene function prediction method using a single expression dataset, we explored three distinct integration approaches to leverage information from multiple transcriptomics studies: (1) simple concatenation of expression datasets (“Full compendium”), (2) aggregation of individual networks (“Averaged networks”) and (3) a novel integration protocol, where individual networks are assigned to predict different GO terms (“Individual networks vHRR”). For the latter, co-expression networks are built from each atlas (Table 1) and the prediction performance for each GO term in each co-expression network is estimated. Subsequently, only the top-ranking networks for a GO term are used to predict genes for that term. Figure 1h,i shows a cartoon of the protocol and a full description is provided in the Experimental Procedures section.

The popular approach of concatenating multiple expression datasets (“Full compendium”) does not globally improve over using a single developmental dataset (simGIC: Single atlas HRR: 0.045, Full compendium HRR: 0.039). While applying MLC on the full compendium, as intended by the authors (Makrodimitris et al. 2020), improves over its application on a single atlas (simGIC: Single atlas vHRR-MLC: 0.046, Full compendium vHRR-MLC: 0.048, relative increase of 3.2%), this integration approach is not able to outperform the best performing single atlas method (Figure 1a,b).

An alternative to concatenation of expression datasets is to build co-expression networks from each atlas, followed by aggregation of individual networks (by averaging edge weights) to obtain a single integrated network. Subsequently, network propagation algorithms can be applied on the integrated network. Given the promising results with a single atlas, we applied both HRR and vHRR on the aggregated network. Surprisingly, allowing co-expression neighborhood size to vary between GO terms, does not improve over a global HRR threshold (simGIC: Averaged networks HRR: 0.050, Averaged networks vHRR: 0.050). While network aggregation outperforms simple concatenation, the benchmark indicates that this integration method is unable to improve overall performance compared to a single developmental atlas (Figure 1a,b).

Conversely, our novel integration approach, Individual networks vHRR, shows to improve over the second-best performing method by 17.4% (simGIC: Individual networks vHRR: 0.060, Single atlas vHRR: 0.051; Figure 1a). While compared to Single atlas vHRR, the mean specificity of individual predictions slightly decreases (0.58 to 0.55; Figure 1f), the mean number of leaf predictions per gene increases by 27.3% (4.1 to 5.2; Mann-Whitney U p-value < 10^−5^; Figure 1e), highlighting its potential to leverage different functional information captured by individual atlases. However, the improved performance comes with an increased computational time (one order of magnitude; Figure 1g), directly proportional to the number of datasets, and should be considered when choosing the appropriate method.

Global prediction measures provide an overall indication of a method’s performance. However, they offer little insight into which functions are correctly predicted, and how predictions compare across methods. Therefore, we explored the top five processes for each method (referred to as ‘top processes’), showing distinct patterns of functional categories and clarifying differences in global performance (Figure S3). The breadth and distinctiveness of a method, relative to all other methods, is measured by the ‘heatmap score’, where a low score indicates good performance (see Experimental Procedures). While most methods can correctly predict top processes outside their own top five, EGAD, and to a lesser extent BLAST scoring, fail to do so. Moreover, EGAD’s top processes are often better predicted by other methods, which is quantified by a high heatmap score (105; Figure S3). Conversely, the top processes (e.g. ‘amino acid transport’) of BLAST scoring (heatmap score: 40), the only method relying on protein sequence similarity rather than gene expression data, are not successfully predicted by expression-based methods. However, when multiple expression datasets are taken into account, some BLAST scoring top processes improve in prediction quality. Furthermore, expression-based methods using a single dataset are overall less likely to correctly predict top processes compared to methods using multiple atlases, highlighting the benefit of including multiple atlases. Out of all expression-based methods, Individual networks vHRR achieves the highest breadth and distinctiveness considering all benchmark GO terms, as indicated by its heatmap score (41).

From this benchmark, we conclude that our newly developed vHRR method for expression-based AFP outperforms first-generation sequence similarity methods, as well as publicly available state-of-the-art expression-based methods. Moreover, we presented a novel integration protocol that improves upon commonly used integration strategies to combine multiple transcriptomics atlases.

### Validation of novel predictions using experimental functional genomics data produces high-confidence annotations for unknown genes

Given that functional annotation of Arabidopsis genes is far from complete, we employed our integrative AFP approach to assign novel functions to both known and unknown genes, using the transcriptomics studies reported in Table 1 and experimental and curated annotations as input data. This procedure predicted a total of 1,196 BP terms for 23,470 genes, of which 9,008 are currently lacking any BP annotation. However, relying solely on co-expression GBA to infer functional annotations is likely to produce many false positive associations (Gillis and Pavlidis 2012). To filter out possible spurious associations and provide additional evidence, we directly validated our functional hypotheses using experimental protein-DNA interaction (PDI) and protein-protein interaction (PPI) data. To facilitate the validation process, PDI and PPI networks are built and overlapped with the expression-based functional predictions (Figure 2a). A gold standard PDI dataset compiled from high-confidence interactions downloaded from PCBase (Chow et al. 2019) and interactions manually curated from literature (see Experimental Procedures), contains 255 transcription factors (TFs) and 21,308 target genes, with a total of 176,460 interactions. This experimental gene regulatory network (GRN) was used to build functional regulons, defined as co-expression modules enriched for specific TF binding events (Kulkarni et al. 2018). If a regulon is enriched for a GO term annotated to the incoming TF, each target gene within that regulon and predicted with that term is considered to be validated by an experimental PDI. This approach adds functional context to TF-target interactions, limiting spurious associations inherent to some experimental methods, such as chromatin immunoprecipitation sequencing (ChIP-Seq). High-confidence PPIs were downloaded from IntAct (Orchard et al. 2013), BioGRID (Oughtred et al. 2021) and the Arabidopsis Interactome (Arabidopsis Interactome Mapping Consortium 2011). In the resulting PPI network, covering 9,290 proteins and 43,626 interactions, experimental and curated annotations are propagated to a protein’s direct neighbors and a functional prediction is considered validated by PPI when it coincides with a propagated annotation (Figure 2a).

**Figure 2.**
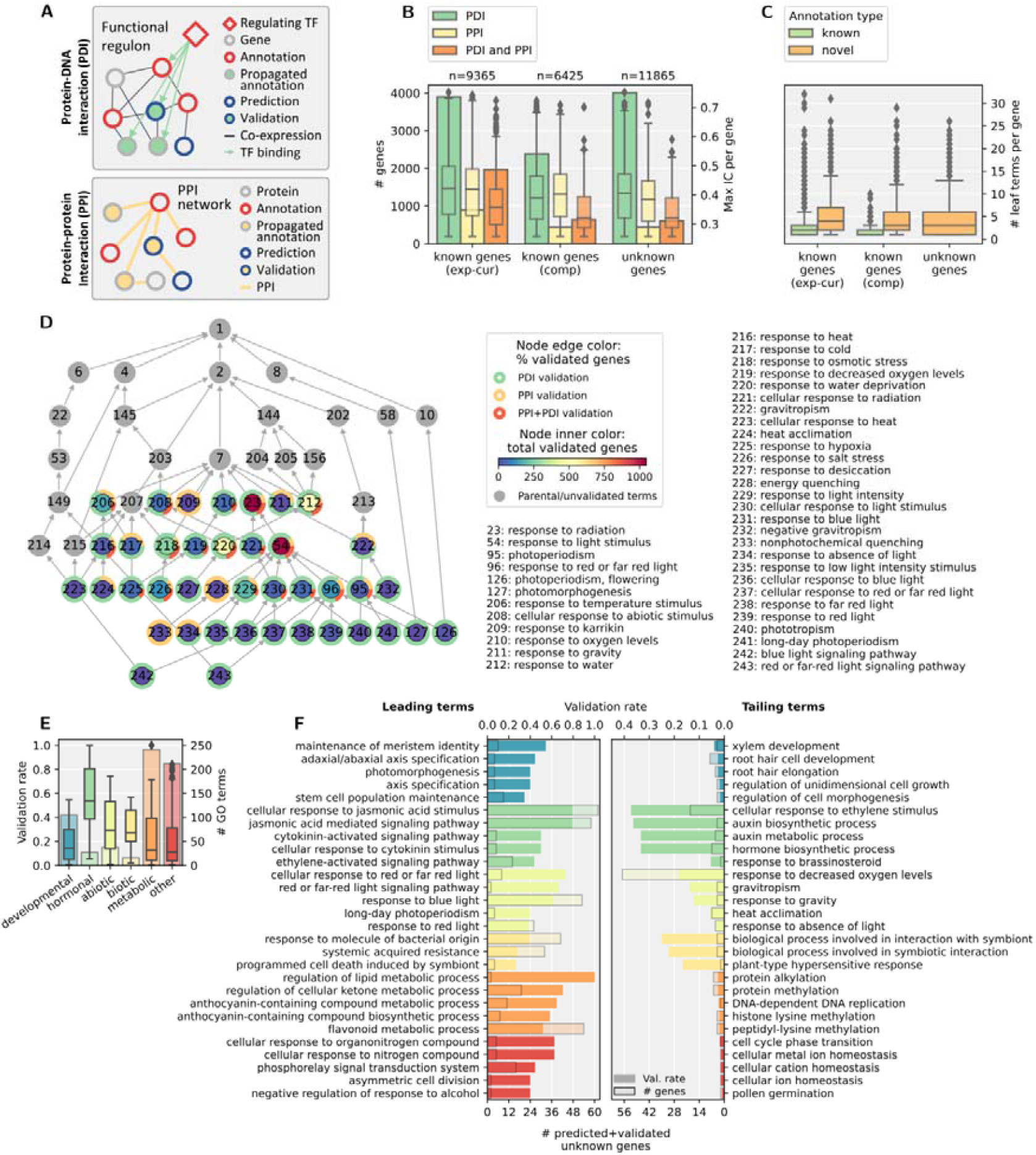
Validation of co-expression-based functional predictions with experimental data. **(A)** Validation protocols using PDI and PPI data. Genes are represented by nodes, while biological interactions are depicted by network edges. Functional regulon: co-expression modules enriched for specific TF binding events; propagated annotation: annotation transferred from binding TF or interacting protein. **(B)** Number of genes with validated functional predictions. Gene counts are depicted by the bar chart, while the boxplots show the distributions of the highest predicted IC per gene for the respective categories. The total number of genes (with or without predictions) per category is shown on top of the chart. While a gene can have multiple predictions, only the gene’s highest confidence (“PDI and PPI” > “PDI” or “PPI”) and most specific (highest IC) prediction is shown. IC: information content, exp-cur: experimental or curated evidence types, comp: computational evidence types. **(C)** Number of known annotations or validated predictions per gene, after removal of parental terms. **(D)** Directed acyclic graph (DAG) representation of the validated functional predictions for abiotic stress biological process terms to unknown genes. Nodes represent GO terms and edges represent “is_a” links in the GO hierarchy. Parental terms are included to extend the DAG towards the root of the tree: “biological process” (GO:0008150, index 1). Node edge colors denote the square root of validation category fractions for the respective term. Table S2 provides a full list of plotted GO terms. **(E-F)** The fraction of validated predictions to all predictions for each GO term (validation rate), grouped into six process categories. Only predictions to unknown genes are taken into account. **(E)** Boxplots show the validation rate distributions, while the bar charts show the number of considered GO terms. **(F)** Validation rate and number of validated predicted genes for the leading/tailing (i.e. top five highest/lowest validation rate) terms with IC>0.5 for each category.

From the set of 175,310 novel co-expression-based functional hypotheses, 65,125 (37.1%) were validated with PPI or PDI, and 8,831 (5.0%) by both data sources, with a combined coverage of 15,239 genes and 744 BP terms (Table S1). Figure 2b depicts annotation (i.e. validated prediction) counts, where for each gene only the prediction with highest confidence and specificity is shown. Genes are separated by the type of known annotation evidence codes, allowing a comparison between known and unknown genes. While predictions to known genes are more often validated than unknown genes (exp-cur: 33.5%, unknown: 25.6%), the specificity of predictions is similar across all three gene categories, indicating that our method does not suffer from assigning more general GO terms to unknown genes. Additionally, the number of leaf (i.e. after removal of parental terms) validated predictions per gene is similar for the three gene categories (mean known (exp-cur): 5.13, mean known (comp): 4.44, mean unknown: 4.23; Figure 2c). In summary, 4,446 and 608 genes without previous BP annotations were predicted and validated by one or two experimental data sources, respectively.

To gain insights on which processes we were able to provide annotations to unknown genes, GO directed acyclic graphs (DAGs) were plotted for four out of six main process categories (developmental, hormonal and abiotic and biotic stresses) and annotated with validated predicted gene counts (Figure 2d, Figure S4). Terms deep in the DAG represent more specific processes compared to their parental terms. Novel validated predictions span all four process categories, for both highly descriptive terms such as “negative regulation of post-embryonic development” (GO:0048581, index 132), as well as more general, yet relevant processes such as “hormone-mediated signaling pathway” (GO:0009755, index 163). The DAGs are instrumental to explore the set of validated predictions for a shortlist of relevant GO terms and pinpoint processes for closer inspection by means of network visualization. Examples of biological processes where a substantial number of unknown genes were predicted and validated are: “root morphogenesis” (GO:0010015, index 83, developmental DAG, PDI: 247 genes, PPI: 9 genes, PDI+PPI: 11 genes), “response to abscisic acid” (GO:0009737, index 191, hormonal DAG, PDI: 387 genes, PPI: 9 genes, PDI+PPI: 55 genes), “response to red or far red light” (GO:0009639, index 96, abiotic DAG, PDI: 115 genes, PPI: 6 genes, PDI+PPI: 9 genes), and “defense response to fungus” (GO:0050832, index 263, biotic DAG, PDI: 296 genes, PPI: 9 genes, PDI+PPI: 8 genes). However, high annotation counts are not expected for all terms, as highly specific processes are likely to involve fewer genes. To investigate if the new annotations of unknown genes do not violate this expectation, the number of genes annotated per GO term was explored (Figure S5). For all process categories, the annotated gene counts follow the trend of the expected distribution (negative correlation between the number of annotated genes and the IC of the GO term), and no outlying terms with high IC and a large number of annotated genes are observed.

Next to absolute prediction and validation counts, the fraction of predictions that were validated for each GO term indicates how well the framework performs in regards to certain types of processes (Figure 2e). Hormone-related processes are most frequently validated, with a median validation rate of 53.8%. Biotic and abiotic stress/response processes are half as likely to be validated (median 27.4% and 29.3%, respectively), and developmental and metabolic processes prove to be the most difficult to predict and validate (median 14.3% and 12.5%, respectively). Despite this, all categories contain specific processes (IC>0.5) with a high validation rate and specific processes with a low validation rate (Figure 2f). For example, several jasmonic acid-, cytokinin- and ethylene-related processes have a validation rate of over 30%, whilst for “response to brassinosteroid” (GO:0009741) only two out of 37 predicted genes were validated. Likewise, meristem-related processes are amongst the developmental processes with the highest percentage of validated predictions, whilst predictions related to root hair formation are less frequently validated by PPI/PDI data sources.

In conclusion, our novel functional annotations contain 6,777 experimentally annotated genes, 3,408 computationally annotated genes and 5,054 unknown genes, covering 774 GO BP terms. Because of the integrative character of the co-expression GBA approach, individual predictions can be traced back to a study-specific network. Thus, gene-centric co-expression modules for predictions originating from the same network can be combined into a single functional module, where validating protein-DNA or protein-protein interactions can be included. In the following paragraphs ‘seed development’ and ‘response to water deprivation’ are inspected closely.

### A developmental multi-omics network links unknown genes to various aspects of seed biology

Seed growth and development play a pivotal role in crop yield, especially for cereal species where seeds are the consumed products. The mechanisms controlling seed development are largely dependent on hormonal signaling cascades and crosstalk, facilitated by abscisic acid (ABA), gibberellic acid (GA), auxin, cytokinin (CK), brassinosteroids (BR) and ethylene (Locascio et al. 2014; Corbineau et al. 2014; Shu et al. 2016). In this study, “seed development” (GO:0048316, index 101, Figure S4) was annotated to 328 (PDI-validated), 39 (PPI-validated) and 18 (PDI+PPI-validated) unknown genes (validation rate: 32.9%). Here, we explored a context-specific multi-omics network, describing co-expression, regulatory and physical interactions, and propose how six previously uncharacterized genes function during seed development (Figure 3). The co-expression network was delineated from a developmental expression atlas, covering several stages of seed development and germination (SRP073212, Table 1). Following the cartoon in Figure 2a, GO:0048316 annotations, predictions, and propagated annotations are highlighted by node color. The color scheme for the six center nodes indicates that the network links these genes with seed development through three complementary data sources. While co-expression analysis can link unknown genes to specific biological processes, insight into how the gene products perform their function is often lacking. The latter can be resolved from physical interactions (PDI and/or PPI), which are indicative of molecular regulatory mechanisms and pathway involvement.

**Figure 3.**
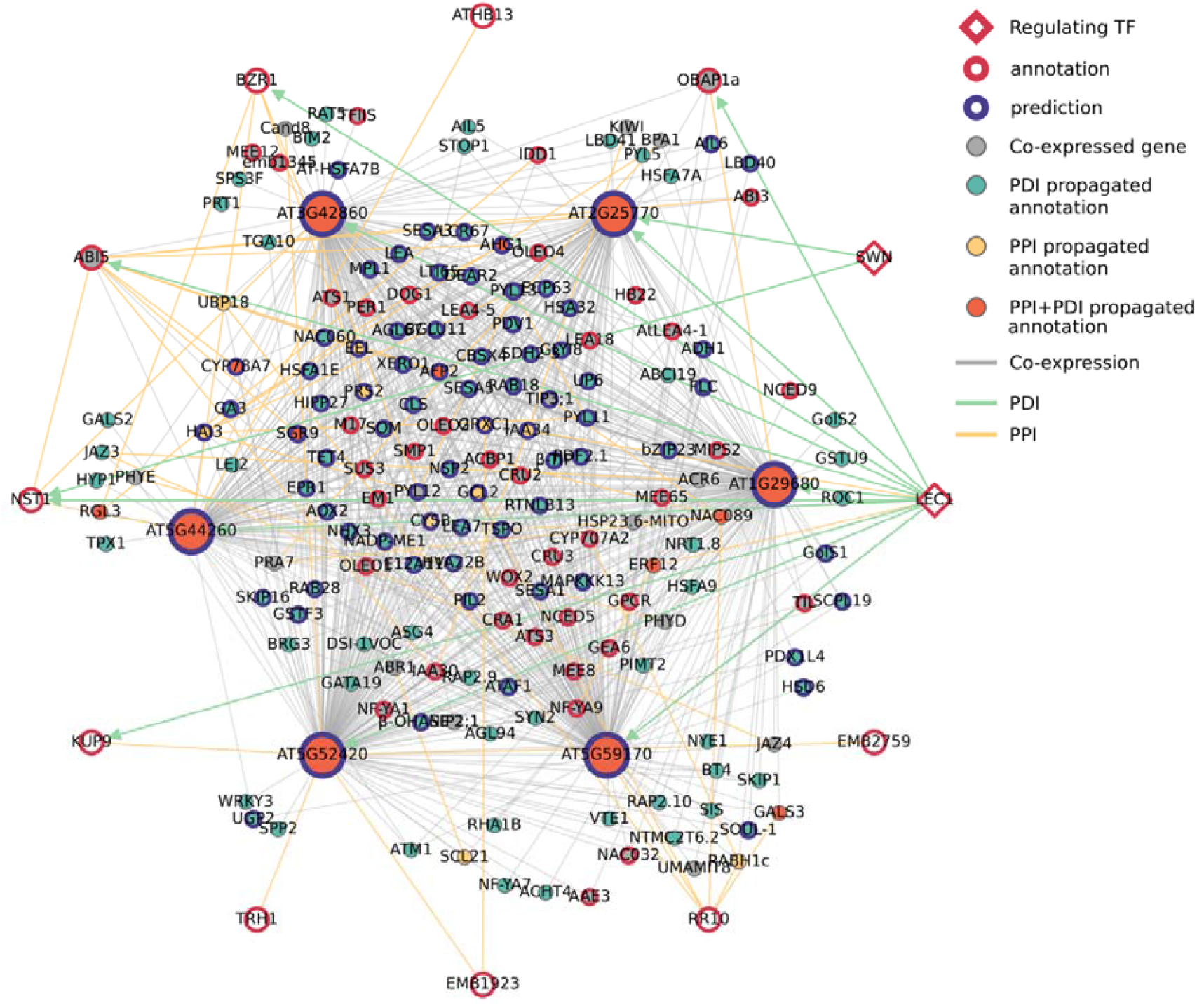
A multi-omics network modeling seed development (GO:0048316). Nodes represent genes and are connected by edges, representing specific gene-gene relationships. Module center genes are represented with an increased node size. The inner cluster shows the optimized vHRR co-expression modules, while nodes plotted on the outer circle are TFs regulating a center gene and its co-expressed genes (PDI), and/or physically interact with a module center gene (PPI). PDI edges are only shown for module center genes and outer circle genes. Co-expression edges are modeled from a developmental expression atlas (SRP075604). Co-expressed genes without an additional edge type (PPI or PDI) or without a gene symbol are omitted for visualization purposes (180 out of 839 co-expressed genes are shown).

**Figure 4.**
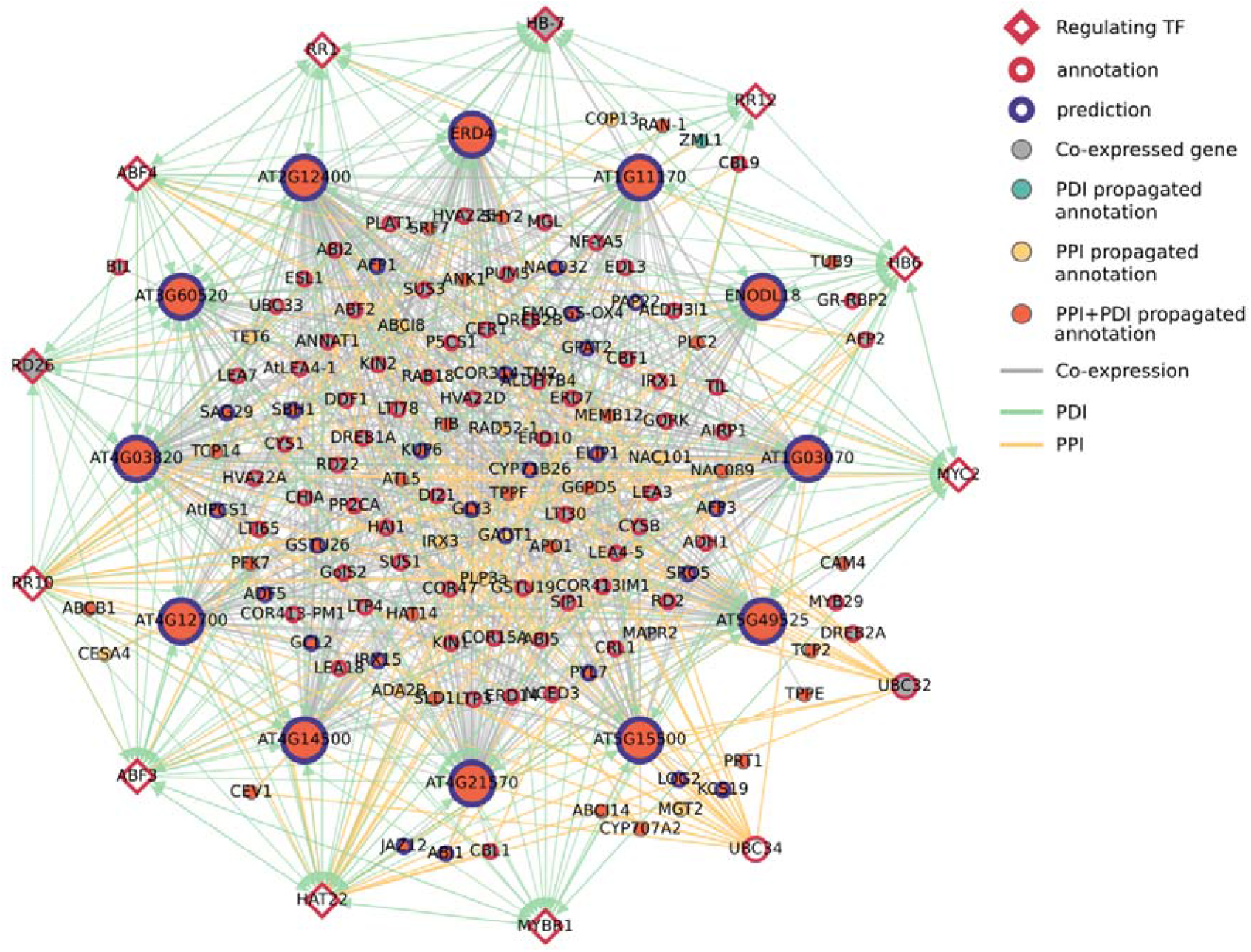
A multi-omics network modeling response to water deprivation (GO:0009414). Nodes represent genes and are connected by edges, representing specific gene-gene relationships. Module center genes are represented with an increased node size. The inner cluster shows the optimized vHRR co-expression modules, while nodes plotted on the outer circle are TFs regulating a center gene and its co-expressed genes (PDI), and/or physically interact with a module center gene (PPI). PDI edges are only shown for module center genes and outer circle genes. Co-expression edges are modeled from a biotic and abiotic stress expression atlas (SRP073212). Only co-expressed genes with a gene symbol and a GO:0009414 annotation, a PPI propagated annotation, or a PPI to a center gene are shown (146 out of 1,667 genes).

For example, the *AT2G25770* co-expression module is regulated by SWINGER (SWN), a core protein in the Polycomb Group (PcG) Repressive Complex 2 (PRC2), which plays a role in controlling the initiation of endosperm development (Wang et al. 2006; Shu et al. 2019). The module is additionally regulated by LEAFY COTYLEDON 1 (LEC1), which is part of the LAFL (LEC1, ABI3, FUS3 and LEC2) transcriptional regulatory network controlling the expression of response genes essential for seed maturation and accumulation of storage compounds (Fatihi et al. 2016). Moreover, the gene product of *AT2G25770* interacts with ABA INSENSITIVE 5 (ABI5), a basic-leucine zipper (bZIP) TF which regulates late embryogenesis-abundant genes and is necessary to maintain quiescence of germinated embryos upon drought stress (Finkelstein and Lynch 2000; Lopez-Molina et al. 2001). Taken together, these results provide strong evidence for a role of *AT2G25770* in seed development, and a suggested function in seed maturation and quiescence maintenance.

All other center genes are similarly regulated by LEC1 and are likely involved in seed maturation. *AT1G29680* interacts with Oil Body-Associated Protein 1a (OBAP1a), which is also a response gene of LEC1 and plays a role in the stability of oil bodies in the embryo (López-Ribera et al. 2014). The AT3G42860 protein interacts with NST1/NAC043, which regulates secondary cell wall thickening in siliques necessary for seed dehiscence, and was linked to seed development due to embryo lethality (Mitsuda and Ohme-Takagi 2008; Meinke et al. 2008). While it is currently unclear how NST1 (and accordingly *AT3G42860*) plays a role in seed development, *AT3G42860* has been significantly linked to seed dormancy in a genome-wide association study (Atwell et al. 2010), further corroborating our biological process annotation. The gene product of *AT5G52420* interacts with THR1/KUP4 and KUP9, two potassium transporters involved in cotyledon development during seed maturation (Tenorio-Berrío et al. 2018). Indeed, AT5G52420 was previously predicted to be a transmembrane protein and the physical interactions with KUP4-9 were identified using the split-ubiquitin yeast two-hybrid assay (Gaudet et al. 2011; Jones et al. 2014). The AT5G59170 protein interacts with RR10, a type-B Response Regulator which mediates the CK transcriptional response and has been shown to negatively regulate seed size (Hill et al. 2013). The gene product of *AT5G44260* interacts with ATHB13, a homeodomain leucine zipper (HD-Zip) TF, regulating response genes during the seed-to-seedling transition, resulting in inhibition of early root growth and affecting cotyledon development upon sucrose signaling (Hanson et al. 2001; Silva et al. 2016). Moreover, AT5G44260 interacts with BZR1, a brassinosteroid (BR)-induced TF which has been shown to negatively regulate ovule and seed number (Huang et al. 2013). These results suggest that *AT5G44260*, expressed in both seed and silique, has various roles in embryo development, as well as a potential role in ovule and seed number establishment.

The presence of several phytohormone-related genes in the co-expression module (e.g. ERF12, IAAs, GA3, ABR1 and JAZs) highlights the roles of ethylene, auxin, GA, ABA and jasmonic acid (JA) signaling pathways during seed development (Locascio et al. 2014; Corbineau et al. 2014; Shu et al. 2016; Ju et al. 2019). In conclusion, this multi-omics network provides strong evidence linking six previously unknown genes to various aspects of seed development. Furthermore, 379 additional unknown genes were annotated with seed development using similar networks (Table S1).

### The leaf water deprivation response is modeled by a drought stress network describing co-expression, regulatory and physical interactions

Understanding the mechanisms which plants use to survive desiccation is crucial in breeding and designing drought-resistant crops. Although drought response and signaling cascades have been studied extensively in Arabidopsis, many links are still missing (Gupta et al. 2020). In this study, 628 unknown genes were predicted with ‘response to water deprivation’ (GO:0009414; index 220; Figure 2d), of which a total of 468 genes were validated (PDI: 424, PPI: 4, PDI+PPI: 40; validation rate: 74.5%). Here, we analyzed a context-specific gene network, describing co-expression, regulatory and physical interactions, and show how twelve unknown genes might function to help accommodate the drought response (Figure 3). The co-expression network was delineated from a transcriptomics dataset sampled from leaf tissues, covering drought stress conditions in conjunction with different biotic stresses (SRP073212, Table 1). The Arabidopsis drought response is regulated by hormonal pathways and crosstalk (Tiwari et al. 2017; Huang et al. 2018; Gupta et al. 2020), of which evidence is present in this combined functional module. ABA (MYC2, HB6, HB7, ABF3, ABF4, RD26, HAT22 and MYBR1), JA (MYC2 and MYBR1), CK (RR proteins) and auxin (MYBR1) related TFs are regulators of the combined co-expression module. All center genes and their respective co-expression modules are regulated by ABA-related TFs, indicating they are ABA-response genes. Indeed, eight out of twelve center genes were also predicted and validated (PDI: *ERD4, AT2G12400, AT4G03820*; PDI+PPI: *AT1G03070, ENODL18, AT4G21570, AT5G15500, AT5G49525*) with ‘response to abscisic acid’ (GO:0009737).

Six out of the twelve center genes (*AT1G03070, ERD4, AT2G12400, AT4G03820, AT4G21570* and *AT5G15500*) interact with the E2 ubiquitin-conjugating enzymes UBC32 and/or UBC34, which have been shown to negatively regulate the ABA-mediated stomatal closure through interaction with the E3 ubiquitin ligase PUB19 at the endoplasmic reticulum (ER) membrane (Ahn et al. 2018). Suppression of these homologous UBC enzymes, alongside a third homolog UBC33, was shown to increase tolerance to drought stress. The reported interactions were identified using the split-ubiquitin yeast two-hybrid assay, indicating that the interacting genes are also membrane proteins (Jones et al. 2014). Thus, these six unknown genes are likely part of the ER-associated protein degradation system and function in the ABA-related drought response. Aside from regulation by ABA-related TFs, *ERD4, AT2G12400, AT4G03820* and *AT4G21570* are regulated by type-B Response Regulator (RR1, RR10 and RR12) proteins, which are thought to negatively regulate the drought response upon CK signaling through restriction of the stomatal aperture (Nguyen et al. 2016). Moreover, *ERD4* has been significantly linked to stomatal aperture in a genome-wide association study (GWAS; Dittberner et al. 2018). Thus, these four genes might help UBC32 and/or UBC34 to enforce CK-mediated stomatal aperture and are novel potential targets in designing drought-tolerant plants.

The remaining six center genes (*ENODL18, AT1G11170, AT3G60520, AT4G12700, AT4G14500, AT5G49525*) are interaction partners of HAT22, an ABA-signaling related HD-Zip TF which inhibits growth and induces senescence upon drought stress through ABA-signaling, but does not play a role in stomatal closure (Liu et al. 2016). All HAT22-interacting center genes, apart from *AT4G12700*, are also regulated by HAT22, and might provide feedback loops in the drought response GRN. Furthermore, the five HAT22-regulated and binding center genes are also regulated by type-B RR-proteins, which, next to the negative regulation of stomatal closure, are thought to inhibit membrane integrity, antioxidant concentration and ABA sensitivity (Nguyen et al. 2016; Huang et al. 2018). Although there have been reports of a link between HAT22 and CK-signaling, the details of the possible interaction are still unknown (Brenner et al. 2005; Liu et al. 2016). Similar to *ERD4*, a single nucleotide polymorphism (SNP) in the *AT3G60520* gene has been identified and significantly linked to stomatal aperture (Dittberner et al. 2018), alluding that *AT3G60520* might also function independently of HAT22 in the CK-mediated regulation of stomatal aperture. Taken together, these five RR-response genes likely play a role at the intersection of HAT22/ABA- and CK-signaling in the drought response, facilitating leaf senescence.

In conclusion, 468 previously uncharacterized genes were predicted and validated with response to water deprivation (Table S1). Moreover, close inspection of a leaf drought response network has shown how twelve genes are linked with the drought response, in particular facilitating stomatal closure/aperture or leaf senescence.

## Discussion

Despite numerous experimental and computational efforts to characterize gene functions in plants, in the model *A. thaliana* still 43% of protein coding genes lack any biological process annotation. In the current study, focused on reducing the plant gene function knowledge gap, the aim was threefold: we set out to (1) develop an improved expression-based AFP approach, applicable in both model and crop species, (2) employ this method to link unknown *A. thaliana* genes to biological processes, and (3) leverage the abundance of largely unexploited experimental functional genomics data to validate the expression-based predictions.

The function prediction performance of our integrative co-expression GBA approach was shown to outperform two state-of-the-art expression-based AFP methods. For this benchmark, a THO validation scheme was adopted, resulting in a highly challenging test set comprising 499 genes. Since the Arabidopsis genome sequence was released in 2000, no experimental evidence was presented linking these genes to biological processes, which were only functionally characterized during the last two years. While the performance metric values (Figure 1b) are much lower than those reported by the third CAFA challenge (Zhou et al. 2019), they are not necessarily indicative of lower performance. Whereas the metrics seem similar, they are not directly comparable. First, CAFA3 Arabidopsis-specific results are reported as gene-centric unweighted Fmax scores, meaning that (1) the specificity of predictions was not taken into account, and (2) for each method, the list of gene-GO predictions, sorted by prediction score, was iterated and every prediction score threshold was evaluated with the test set, yielding an optimal threshold where the corresponding F1-score is maximized (reported metric: Fmax). Conversely, in the current study this threshold was optimized using the training data and the resulting list of predictions was evaluated with the test set only a single time (reported metric: F1). While the latter is almost guaranteed to produce lower values (Fmax ≥ F1), a THO test set is generally not available when predicting gene functions outside of a benchmark experiment, as it would result in a decrease of valuable training data. Second, the annotation test sets across benchmarks are composed of different genes and GO terms, of which the impact is exemplified by the THO evaluation by Hansen et al. (2018), where the reported Fmax values for the BP category are equally low as the F1-scores in the current study. Furthermore, a baseline method from CAFA was included here, enabling a relative comparison across benchmarks, which confirms that metric values cannot be compared directly.

Systematic co-expression analysis using the GBA principle is known to introduce spurious gene-gene and gene-function associations (Gillis and Pavlidis 2012). However, a series of optimized strategies for improving analysis of co-expression networks were reviewed recently by Rao and Dixon (2019) and implemented in the current study. First, we employed a directed approach, enabling a concentrated gene-centric search. Second, the newly presented integration strategy allows the use of ‘condition-dependent’ transcriptome data selection, capturing condition-specific co-expression relationships, while maintaining the broad view of a ‘condition-independent’ approach. Third, the applied correlation metric (PCC-HRR) has been shown to outperform other metrics (Liesecke et al. 2018; Figure 1a,b). Moreover, co-expression modules are optimized on a process-by-process basis, further reducing the risk of false positive functional associations inherent to the systematic application of co-expression GBA.

The presented expression-based AFP approach was designed to optimally leverage the availability of transcriptomics data, making it a valuable stand-alone tool for gene function hypothesis generation in crops. However, high-quality functional annotations are lacking in most crop species (Figure S1b), meaning that the to-be-propagated annotations will likely originate from sequence-based annotation pipelines (Bolger et al. 2018). To explore the potential of such an approach, we applied our method to propagate GO BP annotations derived only from InterPro domain annotations (Blum et al. 2021). Firstly, InterPro domain-derived GO BP annotations only cover 31,9% (8,813; Figure S6a) of all protein coding genes, exemplifying the limitations of sequence-based annotation of biological processes to plant genes (see Introduction). By applying our expression-based AFP method using only InterPro-derived BP annotations, the combined coverage increases to 75.4% (20,851 genes). Moreover, from the set of unknown genes predicted and validated by the current study, 21.7% have a novel annotation which was recovered using only InterPro-derived BP annotations as input for the co-expression analysis (Figure S6b). While these recovered validated predictions tend to be more general, the overall specificity of InterPro-based predictions is comparable to predictions derived from experimental/curated annotations (Figure S6c). Taken together, these results show the potential for formulating high-quality gene function hypotheses using only sequence-based input annotations, and thereby increasing the coverage of annotated genes.

Out of all known experimental, curated and computational annotations (with GO terms considered in this study; see Experimental Procedures), 20.4% (53,765 out of 268,099) were recovered by the co-expression analysis. However, despite the adopted precautions mentioned above, functional associations inferred from co-expression analysis should be interpreted as hypotheses and require independent experimental validation (Rao and Dixon 2019). Here, functional hypotheses were validated using two independent experimental functional genomics data types, enabling functional validation on a scale unmatched by focused reverse genetics experiments. Using this multi-omics approach, we annotated 5,054 (42.6% of 11,865) unknown genes to diverse biological processes, and provided novel and confirmed known functions for 3,408 and 576 (53.0% and 9.0% of 6,425, respectively) computationally annotated genes.

For all processes with newly annotated unknown genes, the fraction of validated co-expression-based functional hypotheses was calculated, providing an indication of which processes were better modeled than others. After categorization of processes, we showed that hormonal processes are most frequently validated; however, all six process categories (developmental, hormonal, abiotic and biotic stress/response, metabolic and other) contain both highly specific processes with a high and low validation percentage. This indicates that our multi-omics framework covers a broad spectrum of highly descriptive terms, but also that some processes are rarely validated. Several explanations can be found for the latter observation: (1) a process can be well modeled on the co-expression level, resulting in high quality functional hypotheses, which fail to be validated due to a lack of experimentally profiled protein-DNA/-protein interactions. Alternatively, several stress-related processes are known to be regulated at the post-transcriptional level by regulation of mRNA processing and/or stability (Floris et al. 2009; Vyse et al. 2020). Such processes are potentially difficult to validate using PDI data, as interactions where the target is a protein coding gene might be rare or non-existing. For example, the response to phosphate starvation is thought to be regulated through microRNA-mediated RNA silencing (Floris et al. 2009; Nguyen et al. 2015). Indeed, the co-expression analysis recovered twenty “cellular response to phosphate starvation” (GO:0016036) experimental annotations and predicted GO:0016036 for 98 unknown genes, indicating that this process is well modeled at the co-expression level. However, none of the predicted genes were validated by PDIs due to absence of any experimental/curated GO:0016036 annotations to TFs in our gold standard PDI dataset. (2) A process is modeled poorly on the co-expression level due to a lack of transcriptional regulation, yet results in low quality functional hypotheses due to spurious gene-gene associations inherent to genome-wide co-expression analysis (Gillis and Pavlidis 2012). (3) A process is modeled poorly on the co-expression level due to the absence of relevant transcriptomics experiments. To conclude, the validation percentage provides a guideline into which processes are best modeled by our multi-omics framework.

The annotations provided by the current study were submitted to TAIR, where they can be explored by the Arabidopsis community to facilitate future research, as novel annotations might fill missing links in developmental- or stress-related pathways. For example, we have uncovered five potential novel players at the intersection of HAT22/ABA- and CK-signaling in the Arabidopsis drought response, an instance of hormonal crosstalk which remains to be fully characterized (Brenner et al. 2005; Liu et al. 2016). Likewise, an in-depth analysis of a seed development network revealed various potential roles for multiple unknown genes during seed maturation. Moreover, a selection of the seed development and desiccation response findings are supported by significant GWAS hits, demonstrating that genes part of quantitative trait locus regions can be further prioritized using our GO BP annotations.

In conclusion, we presented a novel expression-based AFP approach, able to formulate high-quality functional hypotheses starting from experimental and curated functional annotation data. Moreover, when experimental functional annotation data is lacking, we have shown that our AFP method can vastly increase the coverage of sequence-based functional annotation pipelines. Finally, by adopting a multi-omics validation strategy, we linked 5,054 unknown *A. thaliana* genes to biological processes, providing novel insights into a variety of developmental processes and molecular responses.

## Experimental Procedures

### Gene expression data

NCBI SRA (Leinonen et al. 2011) was queried on 09/04/2020 for studies matching the following criteria: RNA-sequencing experiment; presence of biological or technical replicates; wild-type (WT) col-0 accession. The resulting list of study accessions was fed into Curse (Vaneechoutte and Vandepoele 2019) for manual curation. Samples not matching the above criteria were discarded and only studies with twenty or more matching samples were retained. Following curation, Curse provides a Prose package which was downloaded and executed using default settings. In short, Prose uses prefetch and fastq-dump from the SRA toolkit (v2.9.0; SRA-Tools - NCBI) to download sequencing data, FastQC (v0.11.7; Babraham Bioinformatics) to detect adapter sequences, Trimmomatic (v0.38; Bolger et al. 2014) to perform adapter clipping and quality trimming, and Kallisto (v0.44.0; Bray et al. 2016) for expression quantification. Pseudo-alignment of reads was done using the TAIR10 genome assembly (Berardini et al. 2015) and the Araport11 (Cheng et al. 2017) structural annotation. Transcript counts were transformed to gene level TPM values. In post-processing, technical replicates (i.e. equal sample identifier, different run identifier) were averaged, in contrast to biological replicates (i.e. different sample identifier and run identifier). Samples were separated per study, resulting in eighteen distinct expression atlases. Genes are considered expressed when min(TPM) > 2.

### Gene Ontology annotation

*Arabidopsis* gene ontology annotations were downloaded from TAIR10 (Berardini et al. 2015) on two separate occasions (gene_associations.tair.gz; 25 December 2018 and 19 January 2021). Only experimental and curated annotations are taken into account (evidence codes: EXP, IMP, IDA, IPI, IGI, IEP, TAS, NAS, IC). Annotation sets were extended with all parental terms and filtered to only contain BP terms. Using the complete training dataset, the Resnik information content (IC; Resnik 1999) was calculated for each GO term *c* as follows:

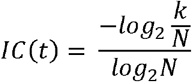

Where *k* is the number of genes annotated with GO term *t* and *N* is the total number of genes with any GO term annotated. All GO annotation sets were filtered to only contain GO terms annotated to [10, 1000[genes.

### Benchmark evaluation setup

A benchmarking study was conducted to evaluate the predictive performance for unknown genes of novel and existing AFP methods. Two measures were taken to avoid evaluation with uninformative GO terms: (1) terms were only included if annotated to 10-1000 genes and (2) evaluation metrics are weighted with the GO IC, so that more specific predictions have a larger impact on a method’s performance. To restrict computational time, the benchmark training dataset (downloaded December 2018) used for the benchmarking study was limited to 500 GO BP terms. In order to not introduce a sampling bias, sets of 500 terms were randomly chosen 10,000 times and the set which showed the highest similarity in IC-distribution with the complete training dataset (minimizing the Kolmogorov-Smirnov statistic) was picked as the benchmark training set. The training process follows a leave-one-out cross-validation (LOOCV) scheme for parameter estimation. In particular, for each method an optimal prediction score threshold is estimated, and predictions below that threshold are disregarded. While LOOCV performance metrics are useful guidelines to see how well training data can be recovered, they do not allow to assess the power of a method to predict functions for unknown genes, unlike a THO validation scheme (Kahanda et al. 2015), which was adopted here. The test set was compiled from annotations downloaded January 2021. Only the training GO terms, extended with parental terms, and only genes not present in the training dataset were retained to obtain the benchmark test set.

Prediction methods included in the benchmark were used to predict gene functions using the benchmark training set, and resulting predictions extended to include parental terms and evaluated using the test set. Two main evaluation metrics were adopted: the gene-centric (GC) weighted F1-score and simGIC (Pesquita et al. 2007):

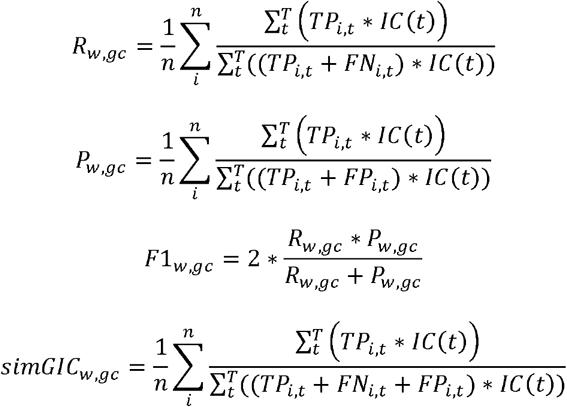

Where *R*_*w,gc*_ is the weighted GC recall, *P*_*w,gc*_ the weighted GC precision, *F1*_*w,gc*_ the weighted GC F1-score, *simGIC*_*w,gc*_ the weighted GC simGIC-score, *n* the number of genes in the benchmark test set, *T* the collection of GO terms in the benchmark test set. *TP*_*i,t*_, *FN*_*i,t*_ and *FP*_*i,t*_ are binary representations of the prediction type of *i-t* (true positive, false negative and false positive respectively) and *IC(t)* the Resnik information content for GO term *t*.

The performance metrics are averaged over all genes, meaning that the coverage of benchmark evaluation genes influences the GC metrics. A high gene coverage is indicative of a general-purpose predictor (Jiang et al. 2016). Likewise, the fraction of covered terms is indicative of a predictor’s ability to model a wide range of plant processes. Depending on the biological context, genes may be involved in different processes. The extent to which this can be captured by each method, is estimated by the number of leaf predictions per gene. The distribution of maximum IC per gene shows the IC of the most specific prediction per gene, also referred to as ‘granularity of functional inference’ (Rhee and Mutwil 2014). In general, more specific predictions, having high IC, are favored over more general ones. Thus, if a method shows good simGIC- and/or F1-scores, but fails to predict highly specific processes, its main performance metrics should be interpreted with caution (Rhee and Mutwil 2014). The computational time for each method is reported as total CPU time.

The top five performing GO terms per method were compared using a heatmap and for each method, a heatmap score was calculated as the number of grey squares in its column summed with the number of colored squares in its top process rows.

### Co-expression network propagation protocol

A novel method was designed to perform GBA gene function prediction using gene expression data. We refer to this protocol as the ‘Single atlas HRR‘ approach, and is implemented in Python3. A fully connected gene co-expression network is delineated where nodes represent genes and edge weights correspond to pair-wise Pearson correlation coefficient (PCC) values. The PCC edge weights are then transformed to highest reciprocal ranks (HRR; Mutwil et al. 2010). A network clustering step is omitted by focusing on gene-centric modules, which requires a HRR threshold and is empirically optimized. For every HRR threshold in a predefined range ([10,400]), gene function predictions are made as follows:

1. For each gene, the GBA process is guided by a GO enrichment analysis of the co-expression module. A leave-one-out cross-validation (LOOCV) is applied by blinding the enrichment step for annotations to the center gene of the co-expression module. The hypergeometric distribution is used to calculate Fisher’s exact p-values, which are interpreted as the prediction score accompanying each gene-GO association.
2. After considering every GO term in the neighborhood of every gene, the set of gene-GO associations is extended by including parental GO terms, projecting the prediction score from the respective child GO terms, unless the parental term was already present in the set with a better prediction score. A Benjamini-Hochberg multiple testing correction (Benjamini and Hochberg 1995) is applied and any gene-GO associations with a corrected p-value larger than the set false discovery rate (FDR) of 0.05 are discarded.
3. The prediction score threshold is optimized by maximizing the weighted F1 (Fmax optimization).

The HRR threshold which results in the highest Fmax is selected as the optimal HRR threshold, and the respective prediction results are selected as the final gene functional predictions.

Alternatively, the optimal HRR threshold can be determined for each GO term individually. For GO terms for which no optimal HRR threshold was found, the mean threshold is selected. Next, predictions are made as described above (steps 1-3), using a variable HRR (vHRR) threshold. This protocol is referred to as ‘Single atlas vHRR’ and is implemented as a Nextflow pipeline (Figure 1h), available at www.github.com/VIB-PSB/vHRR.

### Published AFP methods

The Metric Learning for Co-expression (MLC; Makrodimitris et al. 2020) approach was integrated into our co-expression GBA protocol. The MLC source code was downloaded from GitHub (GitHub - Stamakro/MLC) and converted to Python3. As suggested by the authors, sample weights are learned for each GO term individually. Here, the weighted PCC is used as the initial co-expression metric, from which the HRR network is calculated. Since weights are specific to each GO term, the network calculation, as well as the HRR and prediction score threshold optimization steps are done independently for each term, similar to the Single atlas vHRR protocol described above. The complete procedure was implemented as a Nextflow pipeline.

MLC requires an additional parameter α to be optimized for each GO term. To this intent, an additional optimization loop is included. While parameter α takes any value in the range [0,1], steps of 0.1 were adopted to restrict computational time. For each α, sample weights are learned, followed by network calculation and prediction score threshold optimization. A heuristic is applied to further restrict computational time: values of α which result in a recurring set of weights are disregarded. In particular, if the PCC of a new weight vector with any other previously calculated weight vector exceeds 0.99, the new vector is disregarded and the algorithm moves on to the next value for α.

The EGAD method is implemented in R and available through the Bioconductor project (Ballouz et al. 2017). The following pipeline was constructed:

1. A fully connected gene co-expression network is generated using the provided *build_coexp_network* method, with edge weights representing pairwise Spearman correlation coefficients.
2. Gene functions are predicted using an adapted version of the *predictions* method, which implements a neighbor voting algorithm and returns posterior probabilities for each possible gene-GO combination. Here, an extra step was added to apply a leave-one-out cross-validation, by blinding the neighbor voting algorithm for annotations to the center gene of a co-expression module.
3. The predictions are then processed to find an optimal prediction score threshold based on weighted Fmax, as described previously. The collection of gene-GO associations with a prediction score above the optimized threshold are retained as the final gene functional predictions. Additionally, alternative score thresholds (top 5%, 10% and 50% predictions) were tested.

The BLAST scoring method, as described in the CAFA benchmarks (Radivojac et al. 2013), was included in the benchmark. December 2018 versions of the GOA (Huntley et al. 2015) and SwissProt (Bateman 2019) databases were downloaded. Only annotations to proteins with a sequence available in SwissProt are retained. All annotations were extended to include parental terms, after which the dataset was filtered to only contain the 500 BP terms selected for our benchmark. All corresponding protein sequences were fetched from the SwissProt database, and used to build a BLAST database. Next, the resulting database was queried with all *Arabidopsis* protein sequences using BLASTp, after which self-hits were removed. The BLAST scoring script was downloaded from GitHub (GitHub - Arkatebi/OpenWorld-Problem) and used to score gene-GO associations. The predictions are then processed to find an optimal prediction score threshold based on weighted Fmax, as described above. The collection of gene-GO associations with a prediction score above the optimized threshold are retained as the final gene functional predictions.

### Integrative co-expression analysis using multiple gene expression datasets

A novel integration method was designed, which leverages the specificity of individual gene expression datasets and inferred networks, building on the Single atlas vHRR approach described above. The protocol is implemented as a Nextflow pipeline and follows three steps (Figure 1h,i):

1. The Single atlas vHRR protocol is partly executed for each expression dataset to construct co-expression networks and identify optimal HRR thresholds.
2. For each GO term:
  a. Networks where no optimal HRR threshold was found are disregarded.
  b. From the remaining networks, the mean term-specific Fmax is calculated.
  c. All networks for which the term-specific Fmax exceeds the mean value, are selected for the current GO term.
3. GO-network associations are used to filter enrichment results from each individual network, which are subsequently combined. If the same gene-GO association is made and retained in multiple networks, the prediction with the highest score is selected. After multiple testing correction, the prediction score threshold is optimized using Fmax, resulting in the final gene functional predictions.

Next to the above integration scheme, aggregation of individual networks was tested. Here, a PCC-HRR co-expression network is built from each atlas, after which the networks are aggregated by averaging edge weights: for each gene pair, the connecting edge in the integrated network is weighted with the geometric mean of the corresponding edge weights in the individual networks. Next, the inferred edge weights are re-ranked into the new integrated HRR network. The use of the geometric mean ensures high ranks (i.e. low values) unique to one or few individual networks due, to transient co-expression, to be retained in the integrated network.

### Protein-DNA interaction data and construction of functional regulons

To build an experimental gold standard PDI dataset, all narrow peaks for TFs with two or more replicates were downloaded from PCBase (Chow et al. 2019), except for peaks coming from (Song et al. 2016). The latter were downloaded from the Gene Expression Omnibus (GEO; Barrett et al. 2013; GSE80564) and for each TF, mock and ABA treated samples were interpreted as replicates. Additional ChIP-Seq peaks were downloaded from GEO, for the following TFs: WRKY18, WRKY33, WRKY40 (GSE85922; Birkenbihl et al. 2017); PIF3, ENAP1 (GSE39215; Zhang et al. 2017); ARR1, ARR10, ARR12 (GSE94486; Xie et al. 2018); MYC2, MYC3, STZ, ANAC055 (GSE133408; Zander et al. 2020); CLF, SWN (J. Shu et al. 2019). Next, PDIs were curated from literature for the following TFs: BLJ, JDK (Moreno-Risueno et al. 2015); SCR (Iyer-Pascuzzi et al. 2011); EIN3 (Chang et al. 2013); bZIP1 (Para et al. 2014); GLT1 (Breuer et al. 2012); ARF6 (Oh et al. 2014); REV (Brandt et al. 2012). For all ChIP-Seq data, peaks were annotated to the closest gene, and TGs are only retained when confirmed on the gene level by two or more replicates. Additionally, fourteen Y1H datasets were curated from literature (Brady et al. 2011, Gaudinier et al. 2011, Hussey et al. 2013, Li et al. 2014, Pruneda-Paz et al. 2014, Tian et al. 2014, Jin et al. 2015, Taylor-Teeples et al. 2015, Porco et al. 2016, Sparks et al. 2016, Shani et al. 2017, Gaudinier et al. 2018, Ikeuchi et al. 2018, Smit et al. 2020) and two curated PDI datasets were included (Hussey et al. 2013; Jin et al. 2015). Finally, two more restrictions were applied: TFs must have 10 or more TGs, and a TF must have an experimental or curated GO BP annotation (terms with 10-1000 annotated genes, downloaded January 2021).

Functional regulons were constructed and functionally annotated according to (Kulkarni et al. 2018). In short, a k-nearest-neighbor (kNN) co-expression network was constructed (k=300) for each atlas. Next, all gene-centric kNN co-expression modules were tested for TG enrichment for each TF in the gold standard (adj. p-value < 0.05) to define functional regulons. For each functional regulon, GO terms annotated to the respective TF are tested for enrichment in the regulon (adj. p-value < 0.05), to annotate functional regulons.

### Protein-protein interaction data

IntAct (Orchard et al. 2013) was queried for *A. thaliana* PPIs on 27/10/2020. Likewise, BioGRID (Oughtred et al. 2021) release 4.2.193 was downloaded and queried for *A. thaliana* PPIs. The Proteomics Standards Initiative Molecular interactions (PSI-MI) ontology (Jupp S. et al. 2015) was used to filter interactions based on interaction type (“physical association”: MI:0915 + child terms) and detection method (“biochemical”: MI:0401 + child terms, “biophysical”: MI:0013 + child terms, “protein complementation assay”: MI:0090 + child terms). Interaction were retained which had at least one selected “interaction type” evidence term and at least one selected “interaction detection method” evidence term, and for which Araport11 identifiers were available for both interactors. Additionally, the AI-1, PPIN-1 and NAPPA datasets from the Arabidopsis Interactome were included (Arabidopsis Interactome Mapping Consortium 2011). In the resulting PPI network, where nodes are genes and edges represent interactions, BP terms (annotated to 10-1000 genes, downloaded January 2021) annotated to a node are propagated to the node’s direct neighbors.

## Acknowledgements

We would like to thank Camilla Ferrari for compiling the protein-DNA interaction dataset and Dries Vaneechoutte for his work in the initial development of the co-expression functional inference method. We thank Camilla Ferrari and Indeewari Dissanayake for proofreading the manuscript. We declare no conflict of interest.

## Short legends for supporting information

**Table S1**. Validated predictions and interaction partners

**Table S2**. Directed acyclic graph GO indices

**Figure S1**. Status of plant genomics in 2021.

**Figure S2**. EGAD predictive performance with varying prediction score thresholds.

**Figure S3**. Top performing GO terms for all benchmarked AFP methods.

**Figure S4**. Directed acyclic graphs for BP terms in four process categories with validated functional predictions for unknown genes.

**Figure S5**. Number of predicted and validated unknown genes and GO terms for six process categories.

**Figure S6**. Co-expression analysis using InterPro domain annotations.

