## Supplementary material for "Multi-omics network-based functional annotation of unknown Arabidopsis genes": Figure S

### Supplemental Figures

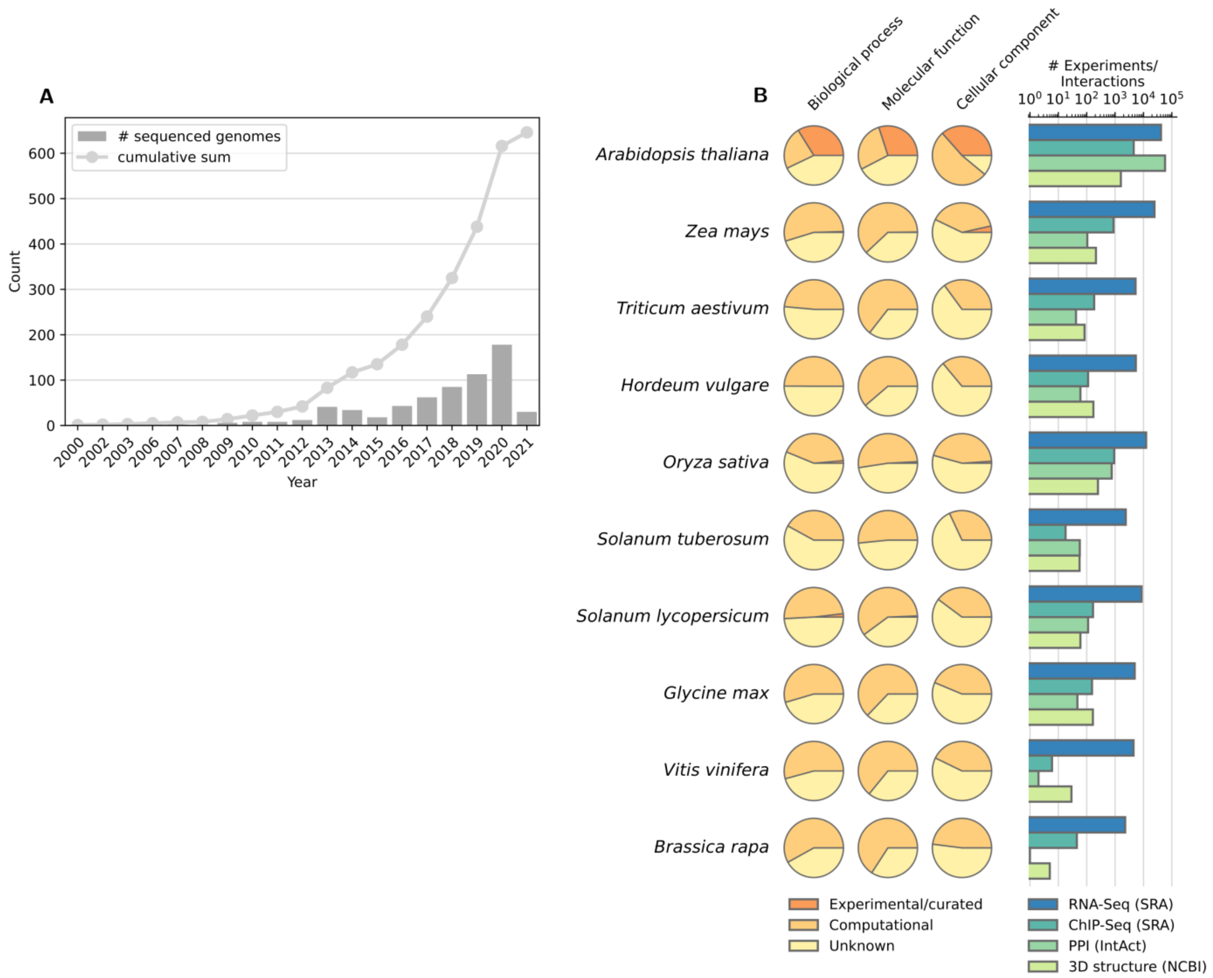


**Figure S1. Status of plant genomics in 2021. (A)** Number of sequenced land plant genomes per year, as reported in the NCBI Genome database (NCBI Resource Coordinators 2018; queried 30/04/2021). Only scaffold or chromosome level assemblies were included. **(B)** Gene function annotation and omics data in plant model and crop species, after Rhee and Mutwil (2014). The pie charts represent, per GO category, the proportion of (un)annotated protein coding genes, according to annotation evidence type. If a gene has both experimental and computational annotations, it is only counted as experimental. For *A. thaliana*, annotations were downloaded from TAIR (Berardini et al. 2015). For all other species, annotations were downloaded from PLAZA4.5 (Van Bel et al. 2018). For *Z. mays*, the PLAZA annotation set was complemented with experimental annotations from MaizeGDB (Portwood et al. 2019). The bar charts show the number of omics experiments or interactions present in public databases per species. For RNA-Seq and ChIP-Seq, the number of experiments present in the SRA database are shown (Leinonen et al. 2011; queried 22/04/2021). For PPI, the number of binary interactions in the IntAct database are shown (Orchard et al. 2013; queried 22/04/2021). For 3D protein structures, the number of structures in the NCBI Structure database are shown (NCBI Resource Coordinators 2018; queried 22/04/2021).

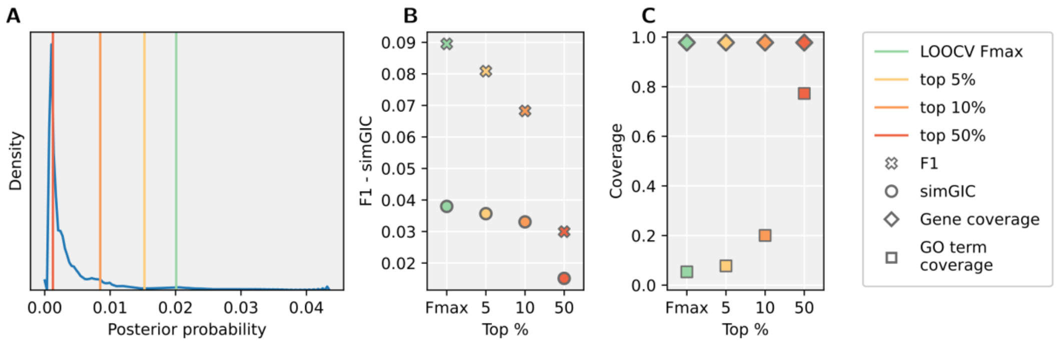


**Figure S2. EGAD predictive performance with varying prediction score thresholds. (A)** Distribution of posterior probability prediction scores of all possible gene-GO combinations. Tested thresholds are highlighted with vertical lines. The leave-one-out cross validation (LOOCV) Fmax threshold corresponds to the top 3.3% predictions. **(B)** Benchmark main performance metrics for the different score thresholds. **(C)** Coverage of the temporal hold out test set genes and GO BP terms.

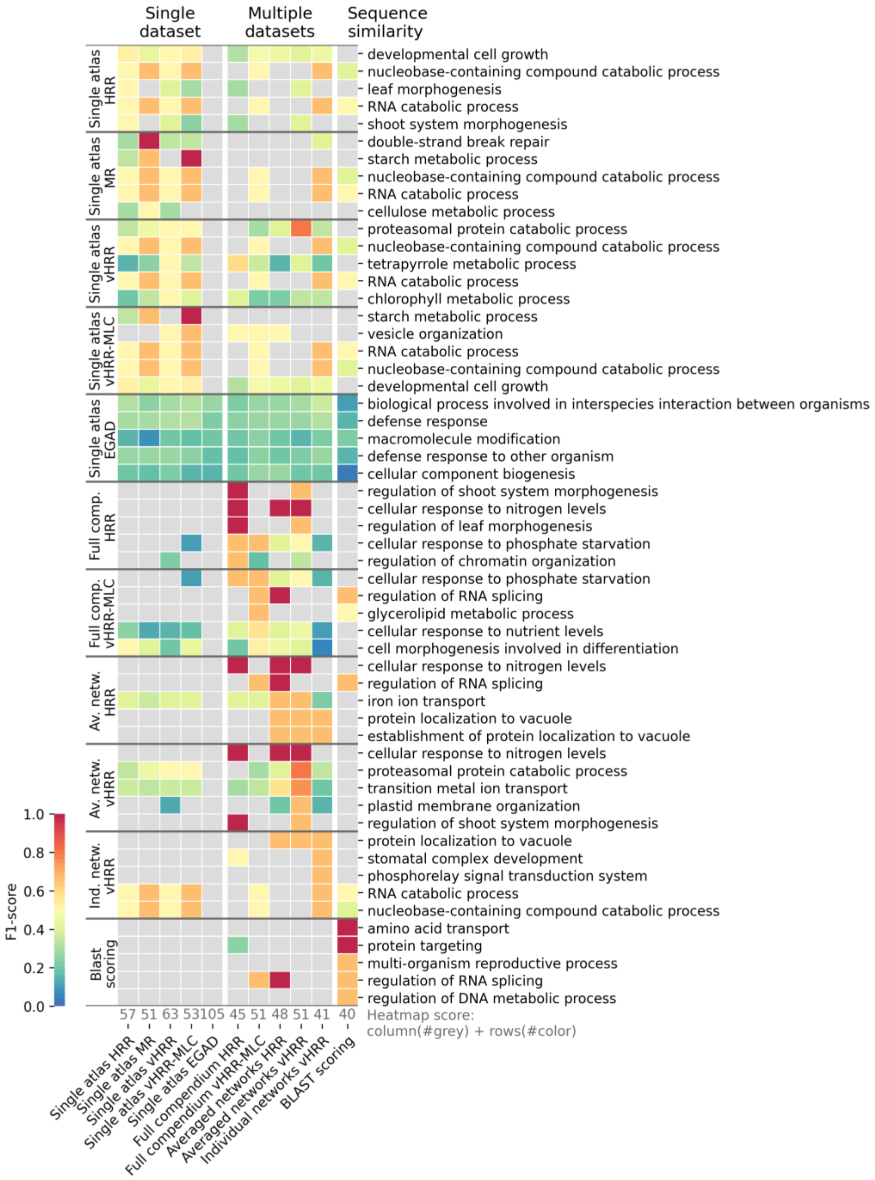


**Figure S3. Top performing GO terms for all benchmarked function prediction methods.** Predicted GO BP terms are ranked based on the term-specific F1-score, and only the top five terms per method are shown. Horizontal lines group the top performing terms per method. A top process can be shared by multiple methods. The “heatmap score” shown in grey equals the number of grey squares in the method’s column summed with the number of colored squares in the method’s top process rows. A low score indicates good performance.

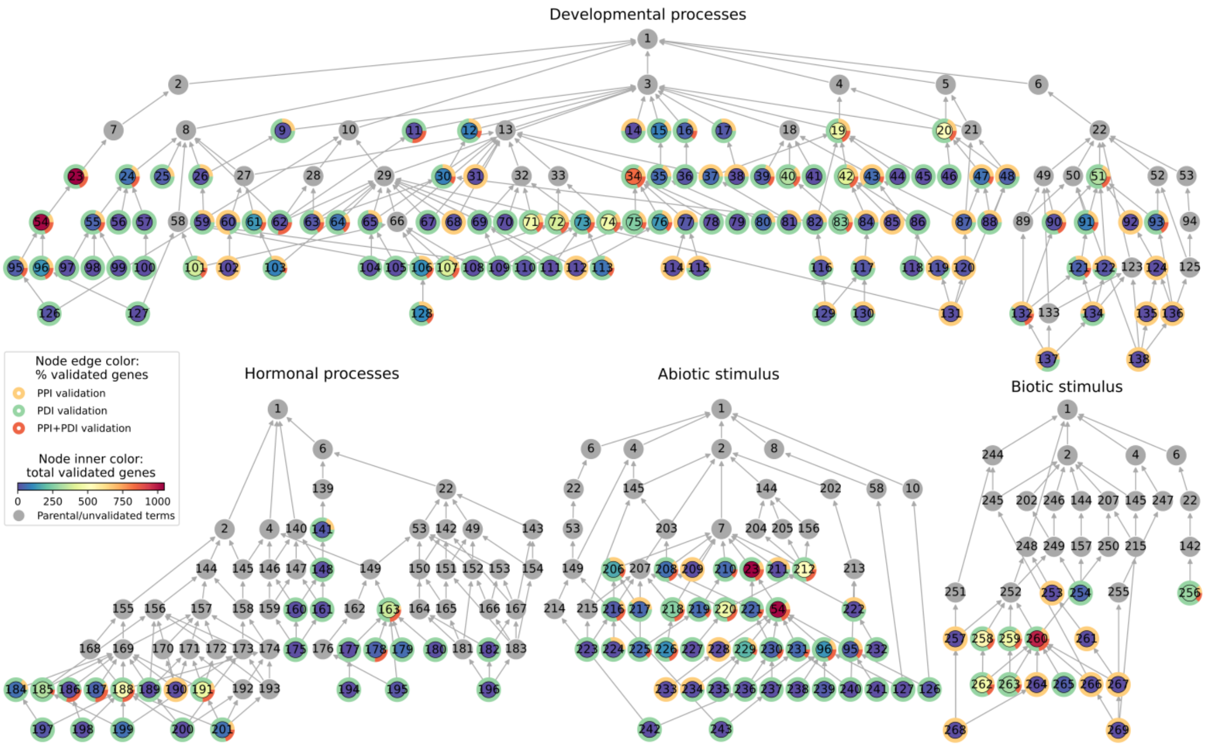


9: maintenance of cell number
11: dormancy process
12: developmental maturation
14: meristem initiation
15: aging
16: anatomical structure formation involved in morphogenesis
17: anatomical structure arrangement
19: cellular developmental process
20: developmental growth
23: response to radiation
24: pattern specification process
25: meristem maintenance
26: stem cell population maintenance
30: anatomical structure maturation
31: root cap development
34: tissue development
35: plant organ senescence
36: plant organ formation
37: cell development
38: meristem structural organization
39: shoot system morphogenesis
40: plant organ morphogenesis
41: post-embryonic plant morphogenesis
42: cell differentiation
43: cell morphogenesis
44: cell fate commitment
45: cell fate specification
46: meristem growth
47: developmental growth involved in morphogenesis
48: developmental cell growth
51: regulation of developmental process
54: response to light stimulus
55: regionalization
56: specification of axis polarity
57: axis specification
59: maintenance of meristem identity
60: gametophyte development
61: embryo development
62: seed dormancy process
63: seed maturation
64: vegetative to reproductive phase transition of meristem
65: floral whorl development
67: anther development
68: pollen tube development
69: floral organ development
70: plant ovule development
71: root development
72: shoot system development
73: phyllome development
74: root system development
75: plant epidermis development
76: meristem development
77: phloem or xylem histogenesis
78: epithelium development
79: epidermis development
80: leaf senescence
81: root hair cell development
82: cell maturation
83: root morphogenesis
84: plant epidermal cell differentiation
85: tracheary element differentiation
86: epithelial cell differentiation
87: unidimensional cell growth
88: root hair elongation
90: negative regulation of developmental process
91: regulation of multicellular organismal development
92: regulation of anatomical structure morphogenesis
93: regulation of developmental growth
95: photoperiodism
96: response to red or far red light
97: adaxial/abaxial pattern specification
98: xylem and phloem pattern formation
99: radial pattern formation
100: adaxial/abaxial axis specification
101: seed development
102: embryo sac development
103: embryo development ending in seed dormancy
104: gynoecium development
105: androecium development
106: reproductive shoot system development
107: fruit development
108: plant-type ovary development
109: stamen development
110: stomatal complex development
111: carpel development
112: embryonic meristem development
113: leaf development
114: phloem development
115: xylem development
116: trichoblast maturation
117: root epidermal cell differentiation
118: epidermal cell differentiation
119: cell morphogenesis involved in differentiation
120: cell tip growth
121: regulation of post-embryonic development
122: regulation of shoot system development
124: regulation of cell morphogenesis
126: photoperiodism, flowering
127: photomorphogenesis
128: flower development
129: root hair cell differentiation
130: trichoblast differentiation
131: pollen tube growth
132: negative regulation of post-embryonic development
134: regulation of flower development
135: regulation of cell morphogenesis involved in differentiation
136: regulation of unidimensional cell growth
137: negative regulation of flower development
138: regulation of pollen tube growth
141: regulation of hormone levels
148: hormone metabolic process
160: auxin metabolic process
161: hormone biosynthetic process
163: hormone-mediated signaling pathway
175: auxin biosynthetic process
177: gibberellin mediated signaling pathway
178: abscisic acid-activated signaling pathway
179: jasmonic acid mediated signaling pathway
180: cytokinin-activated signaling pathway
182: regulation of abscisic acid-activated signaling pathway
184: response to ethylene
185: response to jasmonic acid
186: response to cytokinin
187: response to auxin
188: cellular response to hormone stimulus
189: response to gibberellin
190: response to brassinosteroid
191: response to abscisic acid
194: gibberellic acid mediated signaling pathway
195: ethylene-activated signaling pathway
196: negative regulation of abscisic acid-activated signaling pathway
197: cellular response to ethylene stimulus
198: cellular response to cytokinin stimulus
199: cellular response to jasmonic acid stimulus
200: cellular response to gibberellin stimulus
201: cellular response to abscisic acid stimulus
206: response to temperature stimulus
208: cellular response to abiotic stimulus
209: response to karrikin
210: response to oxygen levels
211: response to gravity
212: response to water
216: response to heat
217: response to cold
218: response to osmotic stress
219: response to decreased oxygen levels
220: response to water deprivation
221: cellular response to radiation
222: gravitropism
223: cellular response to heat
224: heat acclimation
225: response to hypoxia
226: response to salt stress
227: response to desiccation
228: energy quenching
229: response to light intensity
230: cellular response to light stimulus
231: response to blue light
232: negative gravitropism
233: nonphotochemical quenching
234: response to absence of light
235: response to low light intensity stimulus
236: cellular response to blue light
237: cellular response to red or far red light
238: response to far red light
239: response to red light
240: phototropism
241: long-day photoperiodism
242: blue light signaling pathway
243: red or far-red light signaling pathway
253: biological process involved in symbiotic interaction
254: response to molecule of bacterial origin
256: regulation of response to biotic stimulus
257: response to virus
258: response to bacterium
259: response to fungus
260: defense response to other organism
261: biological process involved in interaction with symbiont
262: defense response to bacterium
263: defense response to fungus
264: defense response to symbiont
265: systemic acquired resistance
266: innate immune response
267: programmed cell death induced by symbiont
268: defense response to virus
269: plant-type hypersensitive response

**Figure S4. Directed acyclic graphs (DAGs) for BP terms in four process categories with validated functional predictions for unknown genes.** Nodes represent GO terms and edges represent "is_a" links in the GO hierarchy. GO indices are plotted inside the nodes, with GO descriptions for terms with validated predictions listed below the DAGs. All DAGs shown are extended towards the root of the tree: "biological process" (GO:0008150, index 1). Node edge colors denote the square root of validation category fractions for the respective term. PPI: protein-protein interaction, PDI: protein-DNA interaction. Table S2 provides a full list of plotted GO terms.

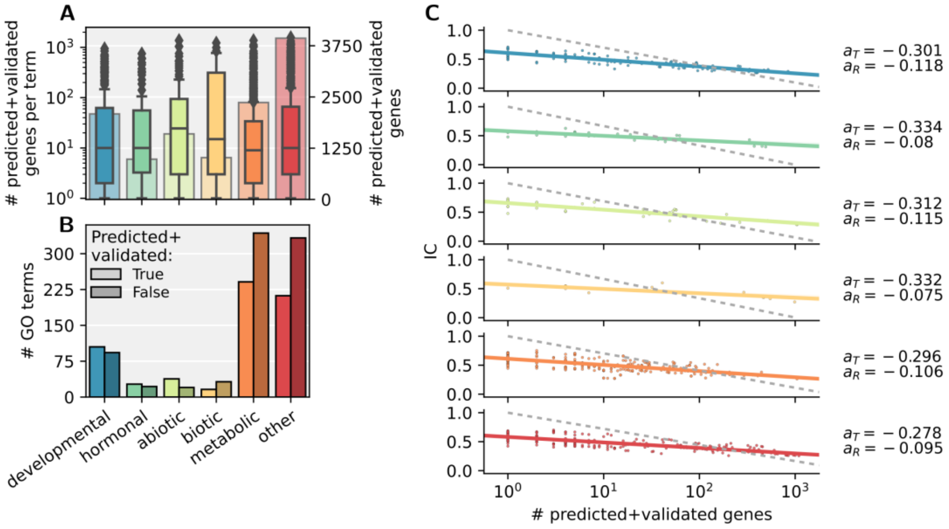


**Figure S5. Number of predicted and validated unknown genes and GO terms for six process categories. (A)** Distributions (boxplots, left axis) and total counts (bar charts, right axis) of the number of predicted and validated unknown genes. **(B)** Number of GO terms with and without newly annotated unknown genes. **(C)** Correlation between the number of newly annotated genes per GO term and the GO information content (IC). Each dot represents a GO term for which at least one unknown gene was predicted and validated. The theoretical IC function (see Methods) when *N* equals the total number of genes within the respective category (as shown in A) is plotted as a grey dashed line. $a_T$ denotes the slope of the theoretical IC function, while $a_R$ denotes the slope of the regression line.

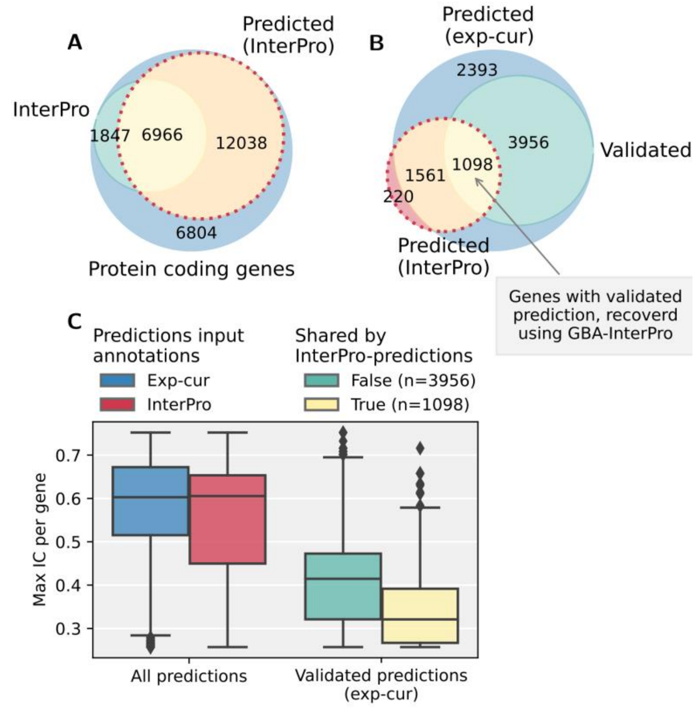


**Figure S6. Co-expression analysis using InterPro domain annotations. (A)** Representation of all protein coding genes, with highlighted fractions of genes with InterPro-derived GO BP annotations and genes with predicted function using our expression-based AFP protocol, using InterPro-derived BP annotations as input. **(B)** Representation of unknown genes, which were functionally predicted using our expression-based AFP protocol, either with experimental/curated BP annotations as input or InterPro-derived BP annotations. **(C)** Information content (IC) of predictions. Left: complete set of predictions from each respective input type. Right: set of predictions using experimental/curated BP annotations as input, validated by additional data sources. InterPro domain annotations were downloaded from PLAZA dicots 4.5 and converted to GO annotations using InterPro2GO.
